# Bayesian estimation on logarithmic scales as an explanation for spatiotemporal interferences with a tendency of deceleration

**DOI:** 10.1101/2020.03.10.986604

**Authors:** Youguo Chen, Andrew Avitt, Minghui Cui, Chunhua Peng

**Author notes:** Corresponding Author: Youguo Chen, School of Psychology, Southwest University, Beibei Chongqing 400715, China.

## Abstract

Spatial and temporal information processing interfere with each other. Kappa effect is a famous spatiotemporal interference, in which the estimated time between two lights increases as an increase of distance between the lights, showing a tendency of deceleration. A classical model attributes the interference to constant speeds and predicts a linear relation, whereas a slowness model attributes the interference to slow speeds and proposes the tendency is the result of the variance of stimuli locations. The present study developed a logarithmic version of the classical model and asserts that the tendency is the result of the Web-Fechner law. These hypotheses were tested in two time discrimination tasks by manipulating the variance of stimuli locations and distance between stimuli. The results demonstrate that estimated time was not modulated by the variance of stimuli locations, and increased as an increase of distance with a tendency of deceleration. The Bayesian model on logarithmic scales made more accurate behavioral predictions than the linear model; the estimated constant speed of the logarithmic Bayesian model was equal to the absolute threshold of speed; the strength of the Kappa effect positively correlated with the variability of time perception. Findings suggest that the interference in the Kappa effect is driven by slow speeds, the strength of the interference is influenced by the variability of time perception, and the tendency of deceleration is the result of the Weber-Fechner law. This Bayesian framework may be useful when applied in the field of time perception and other types of cross-dimensional interferences.

## Introduction

Plenty of tasks require the precise perception of temporal and spatial information, e.g. pilots must land the plane in the right place at the right time. Spatial and temporal representations usually interact with each other in the brain, which has long attracted attention in the fields of psychology and neuroscience [1]. The spatiotemporal interferences offer a window for investigating the nature of representations of time and space [2,3]. The Kappa and the Tau effects are the most famous spatiotemporal interferences. In the simplest experiment, two lights are flashed in sequence to define a distance and a time interval. The time interval seems to increase as an increase of the spatial distance between two lights (Kappa effect)[4,5]. The distance judgments were found to increase as an increase of the time interval between two lights (Tau effect)[6]. Recent research findings show that perceived durations increase with increasing length of lines[2] and area of squares[7]; Correspondingly, a line is perceived to be longer if the line is presented for a longer duration[8].

Some theoretical models focus on the rationale of why different dimensions can interact with each other. A theory of magnitude (ATOM) proposes that time, space and quantity are part of a generalized magnitude system; the inferior parietal cortex is the locus of the common magnitude system[3,9]. A conceptual metaphor theory (CMT) posits that humans understand abstract dimensions (such as time) by mapping them onto their understanding of more concrete dimensions (such as physical space)[2,10,11]. Time is represented on a mental timeline that the past is associated with the left side and the future with the right side[12–14].

Other theoretical models attempt to explain the cross-dimensional interference quantitatively. A classical model of the Kappa effect assumes that observers tend to impute motion to the static lights implicitly, and the model indicates that the imputed motion has a constant speed[5,15–17]. The model quantitatively accounted for the Kappa (or Tau) effect that a temporal (or spatial) judgment in context is a weighted average of the given time interval (or distance) and the expected time (or distance) that would be traversed at a constant speed[18,19]. A slowness model was proposed to explain the cutaneous rabbit illusion, tactile Kappa effect, tactile Tau effect, tactile temporal order judgment and spatial attention effects[20,21]. This model was developed based on a slow speed prior hypothesis. The term slow speed prior comes from the statistical structure of motion that objects tend to be stationary or to move slowly[22–24]. The slowness model combines a slow speed prior with tactile spatiotemporal information to obtain an optimal spatiotemporal perception using Bayes’ rule[20,21].

Recently, a study demonstrated that the classical model can be written as a Bayesian model with an appropriate definition (Fig. 2a)[25]. In the study, two circles flashed from the left to right visual fields in sequence. The time interval and the distance between the two circles were manipulated. Subjects were required to reproduce the time interval between two circles. They found that both the classical model and the slowness model replicated the Kappa effect, but the nonlinear slowness model fits a tendency of deceleration better than the linear classical model, that is, the estimated time increases more slowly with longer distances than that of short distances. The slowness model attributes the tendency of deceleration to the visual acuity in the visual system[20,25]. Visual acuity drops sharply with an increase of eccentricity in the visual system[26]. Due to a poorer visual acuity, the spatial variance of the circle location is larger for long distances than short distances. The slowness model infers that a given physical distance with a larger spatial variance of stimuli locations would be perceived as shorter[27], and that a shorter distance produces a shorter duration perception[20,25]. More evidence should be provided to test the hypothesis of the slowness model on the tendency of deceleration for the visual Kappa effect.

Psychophysical laws are critical for magnitude estimations. Fechner [28] proposed a logarithmic relation between physical magnitudes and the representation by sensory systems based on Weber’s law. Research findings confirm that magnitude estimations are on the logarithmic scales, such as distance reproduction[29–31], numerical quantities[32,33], visual motion perception[23] and time perception[34–36]. The logarithmic function shows a typical tendency of deceleration that the slope of the function decreases with increasing physical distance. This raises our postulate that the tendency of deceleration may be the result of the Web-Fechner law.

Here we proposed a logarithmic version of the classical model (LCM) which incorporates the Weber-Fechner law into the classical model to account for the tendency of deceleration of the Kappa effect (Fig. 2b). The LCM assumes a logarithmic time representation which is based on findings that time discrimination approximately follows the Weber-Fechner law[34–36]. The physical time and the expected time is logarithmically transformed into internal time[23,31,37]. Then the estimated internal time is a weighted average of internal time for physical and expected time, which is similar to the classical model[19,25].

This study presents two experiments that investigate whether the tendency of deceleration is driven by the visual acuity or the Web-Fechner law. The visual acuity hypothesis of the slowness model was tested in Experiment 1 (Fig. 1a). Two sample circles flashed from the left to right visual fields in sequence. The horizontal distance between the two circles and the vertical distance between the circles and the central square were both randomly chosen from three distances (Fig. 1b). The classical model and the LCM predict that the estimated time would keep constant with increasing vertical distance. The slowness model predicts that the estimated time would decrease as the vertical distance increases, because spatial variance of the circle locations increases (or the visual acuity decreases) with increasing vertical distance. Results from the experiment illustrate that the estimated time kept constant with increasing vertical distance, which is in line with the predictions of the classical model and the LCM. Experiment 2 (Fig. 1c) is a continuation of Experiment 1 to test whether the tendency of deceleration is caused by the Web-Fechner law. The distance between the two sample circles was randomly chosen from 1° to 32°, enabling the use of psychophysical functions to estimate the model’s goodness of fit for the tendency of deceleration. Findings show that the LCM rather than the classical model replicated the tendency, which suggests that the tendency of deceleration is driven by the Web-Fechner law.

**Fig. 1.**
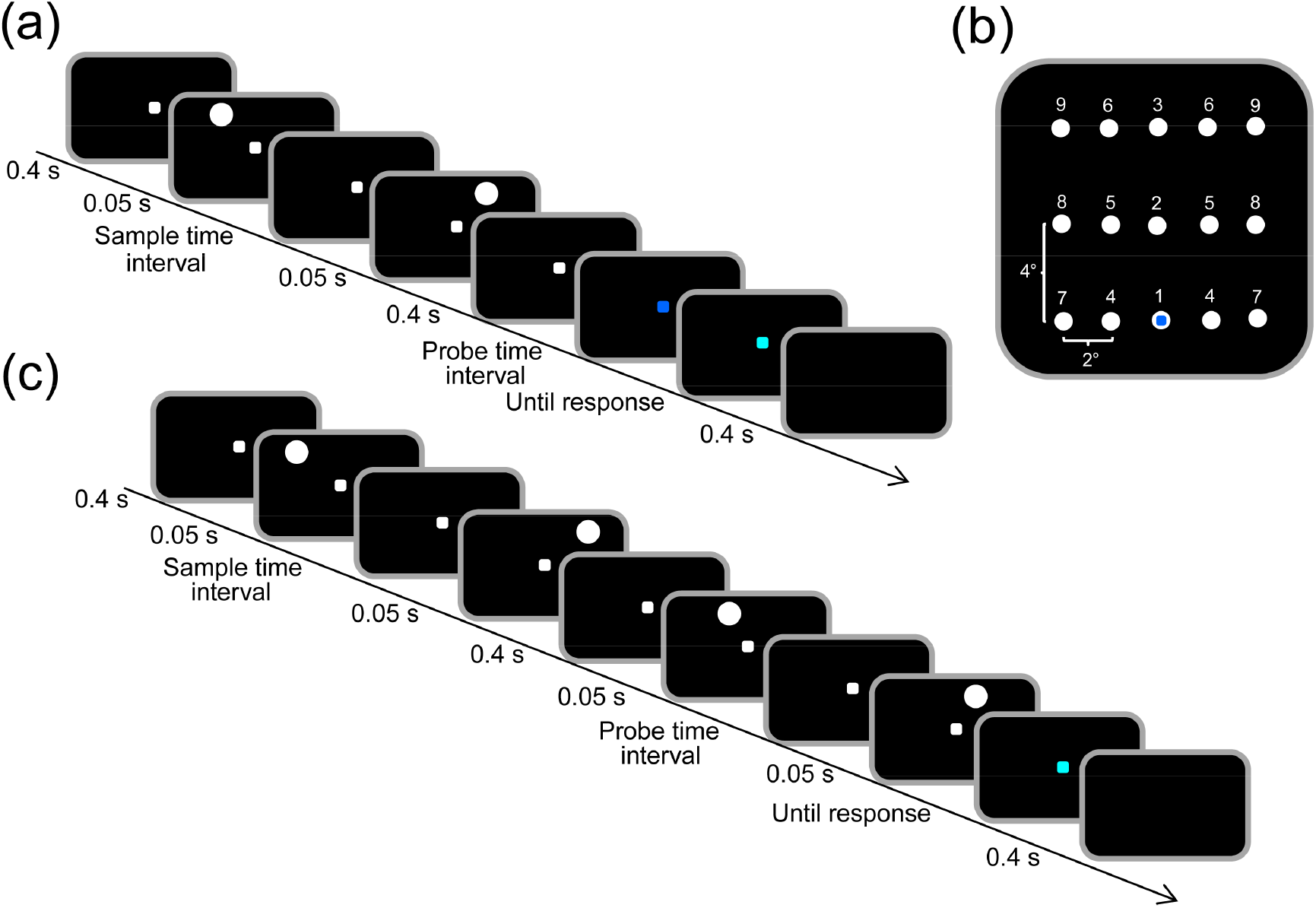
Time discrimination task. (a) The procedure of Experiment 1. The sample was the time interval between two white circles, and the probe was the time interval during the presentation of the blue square. (b) Shows the display positions of circles in Experiment 1. Nine digits represent nine treatments (3 horizontal distances × 3 vertical distances). The fixation square was presented overlapping with the position 1. (c) The procedure of Experiment 2. The sample and the probe were the time intervals between two pair of circles.

## Methods

### Participants

Fourteen participants (6 males, 23 to 27 years of age) took part in Experiment 1, and ten other participants (3 males, 20 to 29 years of age) took part in Experiment 2. All participants had normal or corrected-to-normal visual acuity. Informed written consent was provided by each participant. The project was approved by the Institutional Review Board of Southwest University.

### Stimuli and procedures

The visual stimuli were displayed on a computer screen with a black background. The screen was 27-inch and 60 Hz refresh rate. A white, blue or cyan square was 2 mm in size (0.191°). A white circle was 6 mm in diameter (0.573°). Stimulus presentation was controlled using MATLAB (The MathWorks, Inc., Natick, Massachusetts, USA) and Psychtoolbox 3[38].

### Experiment 1

Experiment 1 was modified from a time reproduction task in the Chen et al. [25] study. We used a time discrimination task with the method of constant stimuli to avoid response bias in the time reproduction task[25,39–41].

Subjects were seated about 60 cm in front of a computer monitor. At the beginning of each trial, a white square appeared in the center of the screen for 0.4 s. They were required to fixate on the central square throughout the trial (Fig. 1a). Two sample circles flashed from the left to right visual fields in sequence, and then the central square turned blue (the probe). The horizontal distance between the two circles and the vertical distance between the circles and the central square were both randomly chosen from three distances (0°, 4°, or 8°). Circle positions were marked by digits from 1 to 9 for nine treatments (3 × 3, Fig. 1b). The sample interval between two circles was 0.5 s. The probe time interval during the presentation of the blue square was randomly chosen from seven durations (0.2 s, 0.3 s, 0.4 s, 0.5 s, 0.6 s, 0.7 s, or 0.8 s). The circles were presented in advance of the blue square in half of the trials, and the blue square was in advance in the other half of trials. Finally, the central square turned cyan as a response signal. We used a two-alternative forced choice (2AFC) experimental protocol, in which subjects were asked to select whether the probe time interval of the blue square was shorter or longer than the sample time interval between the two circles. Subjects pressed “F” or “J” on the keyboard using the index fingers of both hands (“F” denoted shorter, “J” denoted longer). An inter-trial interval of 0.4 s was used.

We used three horizontal distances, three vertical distances and seven probe time intervals, and each treatment consisted of 20 trials, a total of 1260 trials (3 × 3 × 7 × 20) in Experiment 1. Participants would have a short break (about a minute) once they completed 126 trials.

### Experiment 2

Experiment 2 used a time discrimination task with the staircase procedure to save time compared with the method of constant stimuli (Fig. 1c). Two sample circles flashed from the left to right visual fields in sequence. A vertical distance between the circles and the central square was 2.865° (30 mm)[25]. A horizontal distance between circles was randomly chosen from six distances (1°, 2°, 4°, 8°, 16°, or 32°). Sample time intervals between two sample circles were randomly chosen from two durations (0.5 s, or 1 s). Then two probe circles flashed from the left to right visual fields in sequence. The horizontal distance between two probe circles was 1°. Probe time intervals between two probe circles were adjusted according to interleaved adaptive staircase procedures. One staircase started from 0.4 times of the sample time interval (e.g. 0.2 s for the sample time interval of 0.5 s), the other staircase started from 1.6 times of the sample time interval. Staircase procedures consisted of the simple up-down method (one up one down) with a step size of 0.1 times of the sample time interval. The sample circles were presented in advance of the probe circles in half of the trials, and the probe circles were in advance in the other half of the trials. Participants were asked to select whether the time interval between the first pair of circles was shorter or longer than that between the second pair of circles. Participants pressed “F” or “J” on the keyboard using the index fingers of both hands (“F” denoted longer, “J” denoted shorter).

The experiment used two sample time intervals, six horizontal distances. Each treatment consisted of two staircases with 40 trials in each staircase. The 80 trials determined a psychometric function for a treatment[23]. All staircases were randomly interleaved. There were 960 trials (2 × 6 × 2 × 40) in Experiment 2. Participants were permitted a short break (about a minute) once they completed 96 trials.

### Bayesian models

#### Classical model

The classical model assumes that observers tend to impute motion to the stationary stimuli presented successively[5,19,42]. The estimated time *t*_e_ is a weighted average of the actual time *t*_s_ and the expected time *E*(*t*). The expected time is calculated as the ratio of a known distance *l* and a constant velocity *v*_0_[19].

Chen et al. [25] showed that the classical model can be deduced from an appropriately defined Bayesian model (Fig.2a). The model was modified from an ideal observer model for time reproduction[43]. The first stage is a measurement. A sample time interval *t*_s_ is measured as *t*_m_. The distribution of measured time interval *p*(*t*_m_|*t*_s_) is a Gaussian distribution *N*(*t*_s_, *σ*_m_). The mean and standard deviation of measured time interval are *t*_s_ and *σ*_m_ respectively. The second stage is a Bayesian estimator. Participants have a prior belief of a constant speed *v*_0_ of movement. Given that the distance between two flashed stimuli is *l*, the distribution of a prior *p*(*τ*) is a Gaussian distribution *N*(*l*/*v*_0_, *σ*_τ_). The prior *p*(*τ*) and the likelihood *p*(*t*_m_|*τ*) can be integrated into a posterior distribution using Bayes rule, that is, 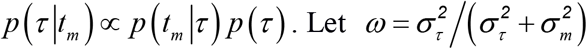. Given a sample interval *t*_s_, the mean of estimated time *t*_e_ is:

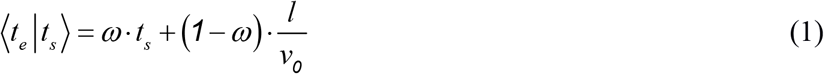

According to the scalar variability of time perception, the standard deviation of a internal time increases linearly with its mean, that is *σ_m_*=*w_m_t_s_*, in which *w_m_* is a Weber fraction[34,35,44,45]. The variance of the estimated time is:

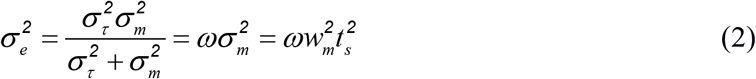

We applied the classical model to fit the psychometric function in the time discrimination task (Fig. 1a). ⟨*t*_*e1*_|*t*_*s1*_⟩ and ⟨*t*_*e2*_|*t*_*s2*_⟩ are means of the estimated time for a sample time interval *t_s1_* and a probe time interval *t_s2_*. *σ_e1_* and *σ_e2_* are standard deviations of the estimated time. In the decision stage, the performance of the time discrimination task is governed by the following equation according to standard signal detection theory[22,46]:

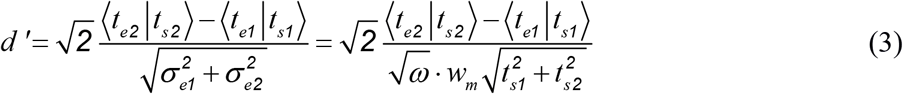

For a range of sample time intervals and probe time intervals, the probability of choosing the probe time interval longer than the sample is determined by:

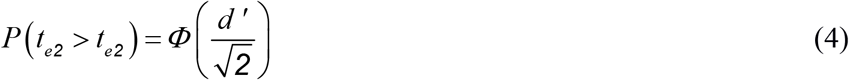

In which Φ denotes the cumulative standard normal distribution function.

At the point of subjective equality (PSE), the estimation of the sample time interval is equal with that of the probe time interval, ⟨*t*_*e1*_|*t*_*s1*_⟩ = ⟨*t*_*e2*_|*t*_*PSE*_⟩, that is:

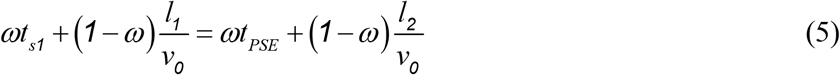

The classical model infers the PSE *t*_*PSE*_ is a linear function of horizontal distance between two sample circles *l_1_*.

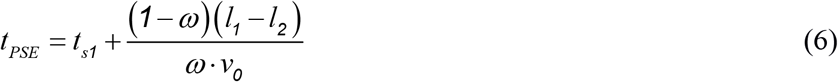

**Fig. 2.**
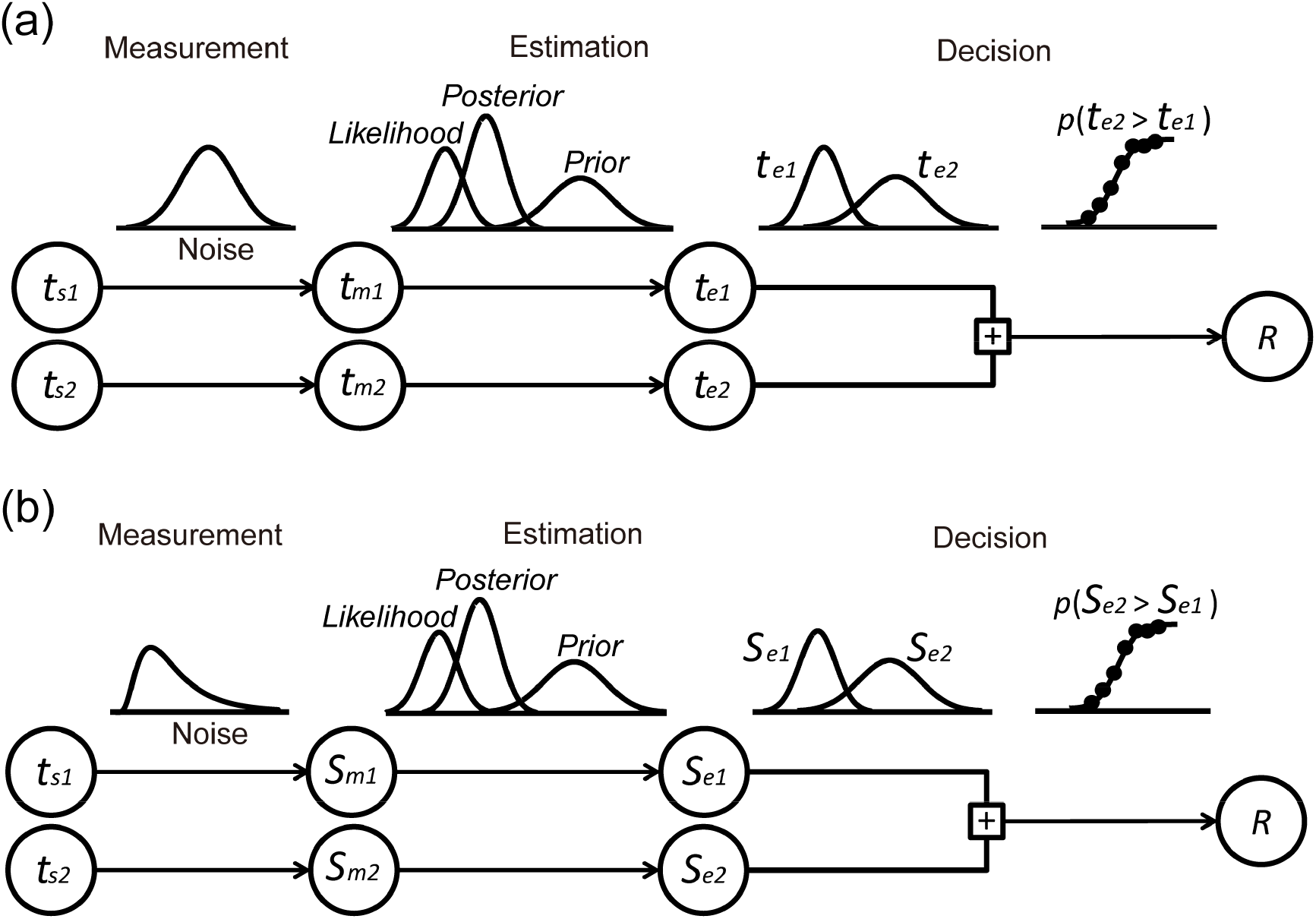
Bayesian models for time discrimination task. On each trial, the observer independently performs an optimal time estimation on sample and probe time intervals and then selects the longer estimate in the decision stage. The responses can be modeled using standard methods from signal detection theory. (a) The classical model assumes a normal likelihood function. (b) A logarithmic version of the classical model (LCM) assumes a log-normal likelihood function which can be logarithmically transformed into a normal distribution.

#### Logarithmic version of the classical model

A logarithmic version of the classical model (LCM) is deduced based on the constant speed prior and the Web-Fechner law (Fig. 2b). Previous studies emphasized the scalar variability in time perception tasks, that is, timing accuracy is relative to the size of the interval being timed. This is consistent with a notion that time discrimination follows the Weber-Fechner law[34,35,44,45]. We used a log-normal likelihood function to model the time interval[23,47], but used a modified logarithmic transformation to account for deviations from the Weber-Fechner law at short time intervals[23,48]. A three-stage Bayesian model (Fig. 2b) was modified from a Bayesian observer model of movements[23].

In the measurement stage, a time interval is measured and logarithmically transformed into an internal time.

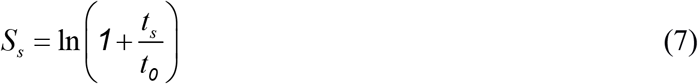

Given a sample time interval *t_s_*, the distribution of an internal sample time interval *S*_m_ is *p*(*S*_m_|*S*_s_), which is a Gaussian distribution *N*(*S*_s_, *σ_s_*_m_).

Participants have a prior belief of a constant speed (*v*_0_) on moving objects. Given the distance between two circles (*l*), the expected time *l/v_0_* is logarithmically transformed into an internal expected time:

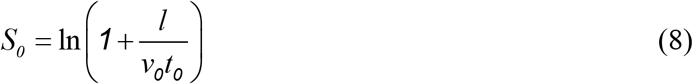

The second stage is a Bayesian estimator. Participants believe a prior *p*(*S_τ_*), which is a Gaussian function *N*(*S*_0_, *σ_sτ_*). The prior distribution is:

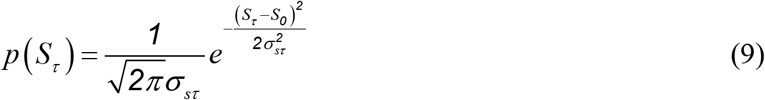

Given an internal expected time *S_τ_*, the likelihood is *p*(*S_m_*|*S_τ_*). The posterior distribution can be computed using Bayes’ rule:

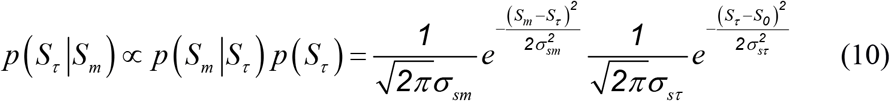

The mean of the posterior distribution is:

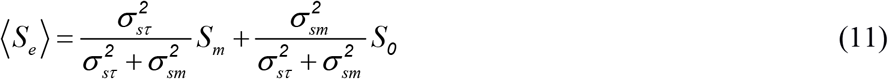

Given a sample time interval *t_s_*, the mean of the internal estimated time is:

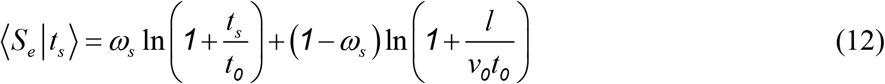

In which 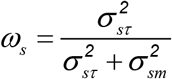, and the variance of the internal estimated time is:

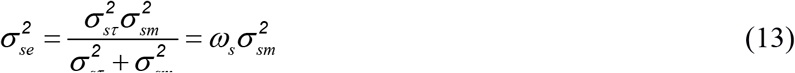

Finally, the internal estimated time *S_e_* is transformed into motor responses *t_e_* using an exponential response model[31,37]:

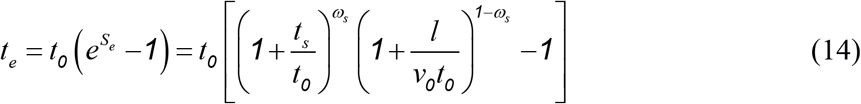

We applied the LCM to fit the psychometric function in the time discrimination task (Fig. 2b). *S_e1_* and *S_e2_* are internal estimated times for the sample and probe time intervals, and *σ_se1_* and *σ_se2_* are corresponding standard deviations. In the decision stage, performance is governed by the signal detection theory:

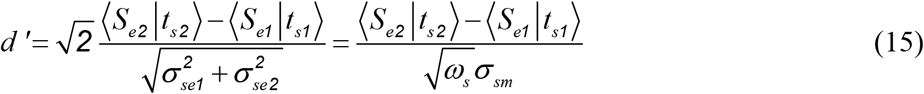

For a range of sample and probe time intervals, the probability of choosing the probe time interval longer than the sample is:

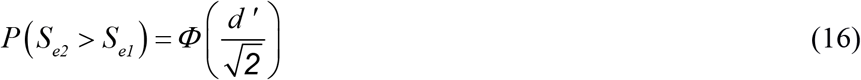

At the point of PSE, the internal estimated times of the sample and that of the probe time intervals are equal, ⟨*S*_*e1*_|*t*_*s1*_⟩ = ⟨*S*_*e2*_|*t*_*PSE*_⟩, that is:

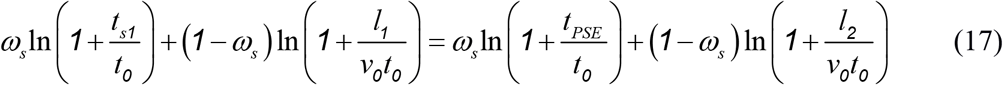

This model infers that the PSE *t_PSE_* is a power function of the horizontal distance of sample stimulus *l_1_*.

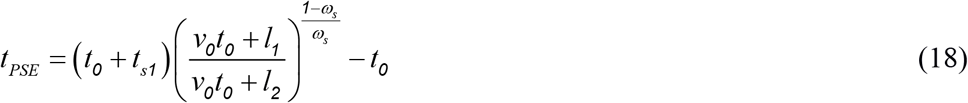

#### Slowness model

The slowness model explains tactile spatiotemporal illusions by integrating the slow speed prior with the likelihood of the neural activities evoked by tactile stimuli[20,21]. Two taps on the skin define a spatial distance *l* and a time interval *t_s_*. *σ*_s_ and *σ_t_* are spatial and temporal standard deviations of neural activities evoked by the taps. The slow speed prior is modeled as a Gaussian distribution *N*(0, *σ*_v_). This model infers that the estimated distance *l_e_* decreases as increasing spatial variability of stimuli locations *σ*_s_:

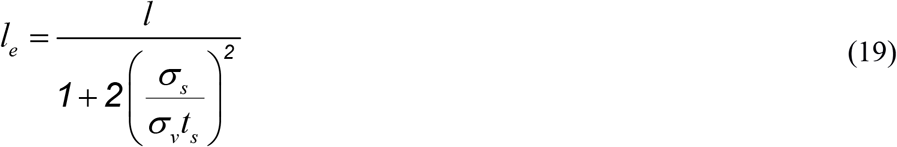

The physical time interval *t_s_* between two taps can be written as a function of estimated time *t_e_*:

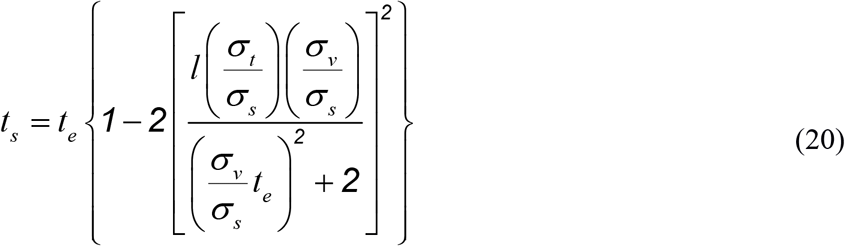

However, the estimated time *t*_e_ can not be written as a function of the sample time interval *t_s_*. The other details of the slowness model can be found in Goldreich’s [20] study, supporting studies by Chen et al.[25].

### Fitting the models to the data

We estimated the PSE (*T_PSE_*) and standard deviation (*σ_PSE_*) of psychometric curves for each treatment and each participant (Fig. 3a and 4a). The maximum-likelihood estimation was used to estimate the best fitting parameters. A response on the i^th^ trial *r_i_* was −1 or +1. Observers estimated the probe time interval as shorter (−1) or longer (+1) than a sample time interval. The probe time interval was *t_s2_*. Let *z* = (*t*_s2_−*T_PSE_*)/*σ_PSE_* and *θ* = (*T_PSE_*, *σ_PSE_*). A probability for the response on the i^th^ trial was calculated using a cumulative standard normal distribution function Φ(*z*):

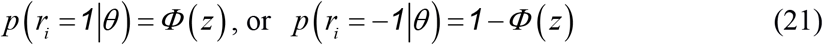

**Fig. 3.**
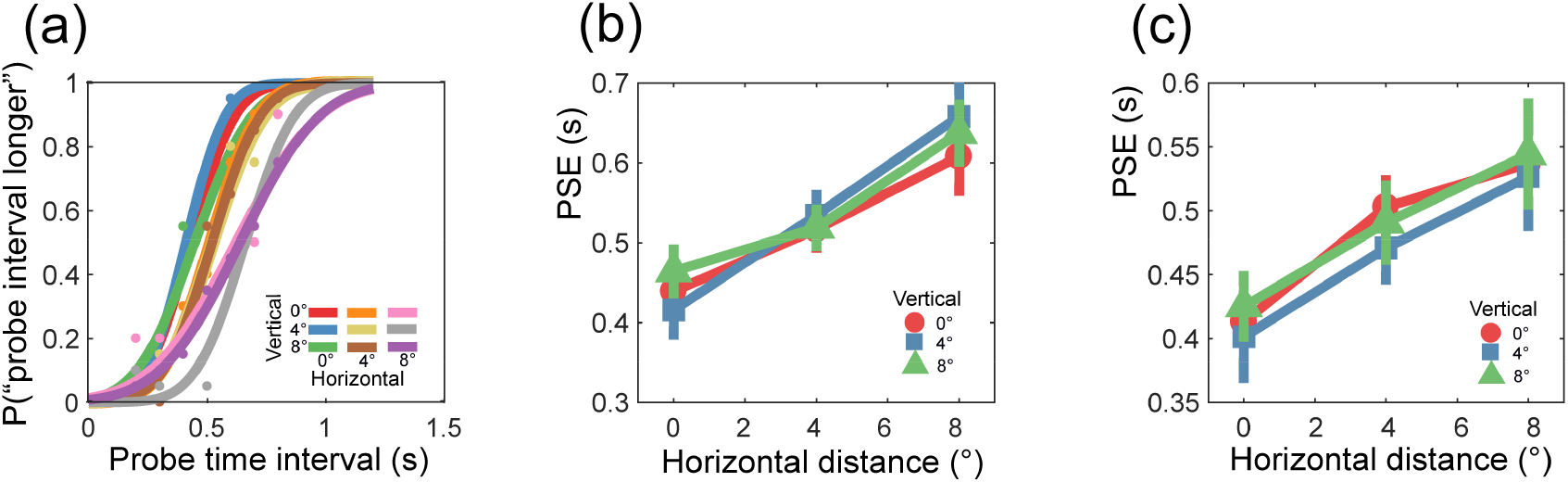
Behavioral responses for Experiment 1. (a) Responses and corresponding psychometric curves of a representative participant. (b) Point of subjective equality (PSE) of a representative participant. The error bars indicated one standard error of mean calculated based on 100 bootstrapped samples of the data. (c) Mean PSE of all participants. The error bars indicated one standard error of the mean across all participants.

**Fig. 4.**
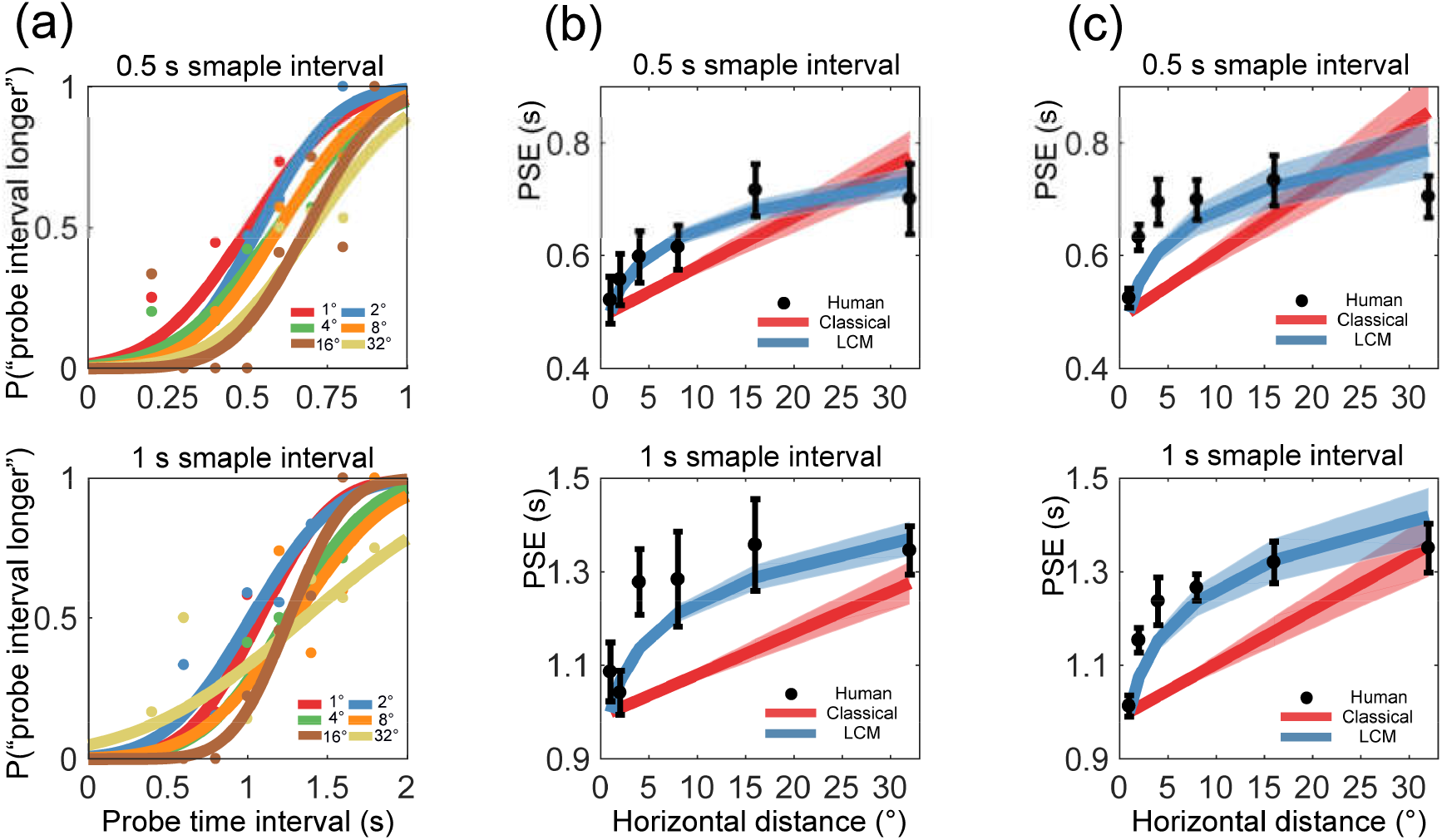
Behavior responses and model fitting for Experiment 2. (a) Responses and corresponding psychometric curves of a representative participant. (b) Point of subjective equality (PSE) of a representative participant, the classical model, and the logarithmic version of the classical model (LCM). The error bars and shadows indicate one standard error of mean calculated based on 100 bootstrapped samples of the data. (c) Mean PSE of all participants and Bayesian models. The error bars and shadows indicated one standard error of the mean across all participants.

Responses were assumed to be independent across trials. The joint conditional probability of the responses across all n trials could be expressed as follows:

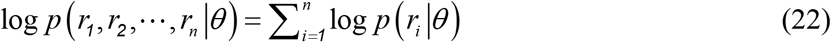

The parameters (*T*_*PSE*_, *σ*_*PSE*_) were found for each treatment and each participant by maximizing the likelihood using the fminsearch command in MATLAB.

To fit the classical model, *d’* was calculated by Equation 3 for each trial. Let 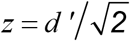, the parameters *θ* = (*ω*, *w*, *v_0_*) were found by Equation 22 for each participant by maximizing the likelihood. Similarly, for the LCM model, *d’* was calculated by Equation 15 for each trial. Let 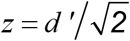, the parameters *θ* = (*ω_s_*, *σ_sm_*, *v_0_*, *t_0_*) were found by Equation 22 for each participant by maximizing the likelihood.

Akaike information criterion (AIC) was adopted to evaluate the goodness of model fitting. AIC was computed for the classical model and the LCM. AIC accounts for overfitting by penalizing models with greater number of parameters[49]. *k* is the number of parameters of the model, *θ* is the best fitting parameters of the model. AIC is calculated as follows, 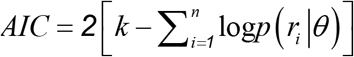. AIC difference (Δ) was obtained by subtracting AIC of the LCM from that of the classical model.

The bootstrap method was used to estimate standard errors of parameters and PSE for the individual participant (Table 1, Fig. 3b and 4b). We resampled data with replacement and repeated this resampling 100 times[23,50,51].

**Table 1.** Parameter estimates from model fits (mean ± SEM).

|  | Classical model |  |  | Logarithmic version of the classical model (LCM) |  |  |  |
| --- | --- | --- | --- | --- | --- | --- | --- |
| | $\omega$ | $v_0$ | $w_m$ | $\omega_s$ | $\sigma_{sm}$ | $v_0$ | $t_0$ |
| S1 | 0.997 ± 0.002 | 0.211 ± 0.158 | 0.745 ± 0.093 | 0.920 ± 0.021 | 0.292 ± 0.089 | 0.085 ± 0.218 | 0.664 ± 0.575 |
| S2 | 0.997 ± 0.001 | 0.310 ± 0.069 | 0.326 ± 0.034 | 0.982 ± 0.006 | 0.051 ± 0.016 | 0.007 ± 0.048 | 3.213 ± 0.947 |
| S3 | 0.998 ± 0.001 | 0.299 ± 0.071 | 0.321 ± 0.026 | 0.948 ± 0.014 | 0.183 ± 0.052 | 0.109 ± 0.118 | 0.486 ± 0.502 |
| S4 | 0.998 ± 0.001 | 0.253 ± 0.047 | 0.455 ± 0.052 | 0.929 ± 0.014 | 0.239 ± 0.053 | 0.111 ± 0.103 | 0.338 ± 0.330 |
| S5 | 0.995 ± 0.004 | 0.205 ± 0.151 | 0.876 ± 0.075 | 0.896 ± 0.040 | 0.293 ± 0.124 | 0.027 ± 0.055 | 1.091 ± 0.968 |
| S6 | 0.998 ± 0.001 | 0.275 ± 0.063 | 0.381 ± 0.041 | 0.939 ± 0.012 | 0.181 ± 0.037 | 0.101 ± 0.106 | 0.343 ± 0.301 |
| S7 | 0.998 ± 0.001 | 0.247 ± 0.050 | 0.440 ± 0.049 | 0.934 ± 0.012 | 0.230 ± 0.047 | 0.106 ± 0.116 | 0.383 ± 0.249 |
| S8 | 0.999 ± 0.0005 | 0.245 ± 0.055 | 0.489 ± 0.061 | 0.936 ± 0.012 | 0.376 ± 0.056 | 0.149 ± 0.107 | 0.095 ± 0.083 |
| S9 | 0.996 ± 0.002 | 0.158 ± 0.085 | 0.672 ± 0.074 | 0.958 ± 0.017 | 0.094 ± 0.042 | 0.036 ± 0.142 | 3.180 ± 1.043 |
| S10 | 0.999 ± 0.0004 | 0.295 ± 0.070 | 0.357 ± 0.039 | 0.956 ± 0.009 | 0.255 ± 0.046 | 0.149 ± 0.097 | 0.182 ± 0.226 |
| All | 0.998 ± 0.0004 | 0.250 ± 0.014 | 0.506 ± 0.057 | 0.940 ± 0.006 | 0.219 ± 0.025 | 0.088 ± 0.015 | 0.997 ± 0.316 |

The standard error of the mean (SEM) was obtained based on 100 bootstrapped samples of the data.

### Statistical analysis

To test whether the variance of the stimuli locations is an influencing factor for the Kappa effect, a two-way repeated measures analysis of variance (ANOVA) was conducted on the PSE for all participants in Experiment 1. The ANOVA factors were Horizontal distance (0°, 4°, 8°) and Vertical distance (0°, 4°, 8°). The Greenhouse–Geisser correction was used to correct for any violations of sphericity[52], and the partial eta squared (*η*_p_²) was used to estimate the ANOVA effect size[53].

## Results

### Experiment 1

Experiment 1 examined whether the variance of stimuli locations modulate the estimated time for the Kappa effect. Two circles were presented in sequence at different locations in Experiment 1 (Fig. 1a and 1b). This allows us to characterize estimated time as a function of the distance between circles and variance of stimuli locations.

The PSE was estimated for nine stimuli locations (3 horizontal distances × 3 vertical distances). For a representative observer, nine psychometric curves clustered into three groups (Fig. 3a). The first group included 0° horizontal distance with three vertical distances (0°, 4°, and 8°), the second group included 4° horizontal distance with three vertical distances, and the third group included 8° horizontal distance with three vertical distances. The PSE increased as an increase of horizontal distances, whereas PSE did not decrease with an increase of vertical distances both for the representative participant (Fig. 3b) and for all participants (Fig. 3c).

The repeated measures ANOVA revealed a significant main effect of Horizontal distance [*F*(2, 26) = 7.655, *p* < 0.05, *η*_p_^2^ = 0.371]. The PSE was significantly smaller in the 0° (mean ± SEM: 0.413 ± 0.026 s) than that in the 4° (0.488 ± 0.018 s) (*p* < 0.05), but the difference between 4° and 8° (0.536 ± 0.034 s) was not significant (*p* > 0.05). The results are consistent with the tendency of deceleration that the estimated time increases more slowly with longer distances than that of shorter distances[25]. The main effect of Vertical distance was not significant [*F*(2, 26) = 2.216, *p* > 0.05, *η*_p_^2^ = 0.146], and the interaction of Horizontal and Vertical distances was not significant [*F*(4, 52) = 0.327, *p* > 0.05, *η*_p_^2^ = 0.025]. The results indicate that variance of stimuli locations is not an influencing factor for the Kappa effect.

### Experiment 2

Experiment 2 supports the assertion that the Weber-Fechner law is critical for the tendency of deceleration in the Kappa effect. A 2-FAC time discrimination task was used to characterize the estimated time as a function of the sample time intervals (0.5 s, 1 s) and horizontal distances between two circles (1°, 2°, 4°, 8°, 16°, 32°) (Fig. 1c). We then fitted the classical model and the LCM to the data, to ascertain which model best explained the data.

The PSE and corresponding standard deviation was estimated for 12 treatments (2 sample time intervals × 6 horizontal distances) for each participant. For a representative observer, the psychometric curves of data moved from left to right as with the increase of the horizontal distance between two circles (Fig. 4a). The results are consistent with the results that the PSE increased with increasing distance between the two circles (Fig. 4b). The tendency of deceleration was observed both for the representative observer (Fig. 4b) and at a group level (Fig. 4c).

A paired sample t-test revealed that the standard deviation of 0.5 s sample time interval (mean ± SEM: 0.265 ± 0.025 s) was significantly smaller than that of 1 s sample time interval (0.423 ± 0.031 s) [*t*(9) = −5.611, *p* < 0.001, Cohen’s *d* = −1.774]. This result is in line with the scalar variability that a longer time interval sample accompanies larger uncertainty[34,35,44,45].

When the models fit to the data, the LCM seems to qualitatively explain human data better than the classical model (Fig. 4b and 4c). The classical model predicts the PSE as a linear function of distance (Equation 6). For the best-fitting parameters (Table 1), *ω* is close to 1, *v_0_* is about 0.3 °/s, and *w_m_* is about 0.5. The LCM predicts the tendency of deceleration that the PSE is a power function of the distance between two circles (Equation 18). For the best-fitting parameters (Table 1), *ω_s_* is about 0.9, *σ_sm_* is about 0.2 s, *v_0_* is about 0.1 °/s, and *t_0_* is about 1 s.

We further explored the individual difference of the Kappa effect based on the LCM. We defined 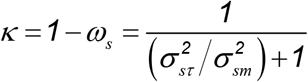 as an index of the strength of the Kappa effect. A larger *κ* indicates a faster increase of estimated time as an increase of the distance *l* between two circles (Equation 12). *σ_sm_* is an index of variability of time perception. We found a significant positive correlation between *κ* and *σ_sm_* (*r* = 0.719, *p* < 0.05, Fig. 5). The result indicates a close relationship between variability of time perception and the strength of the Kappa effect.

**Fig. 5.**
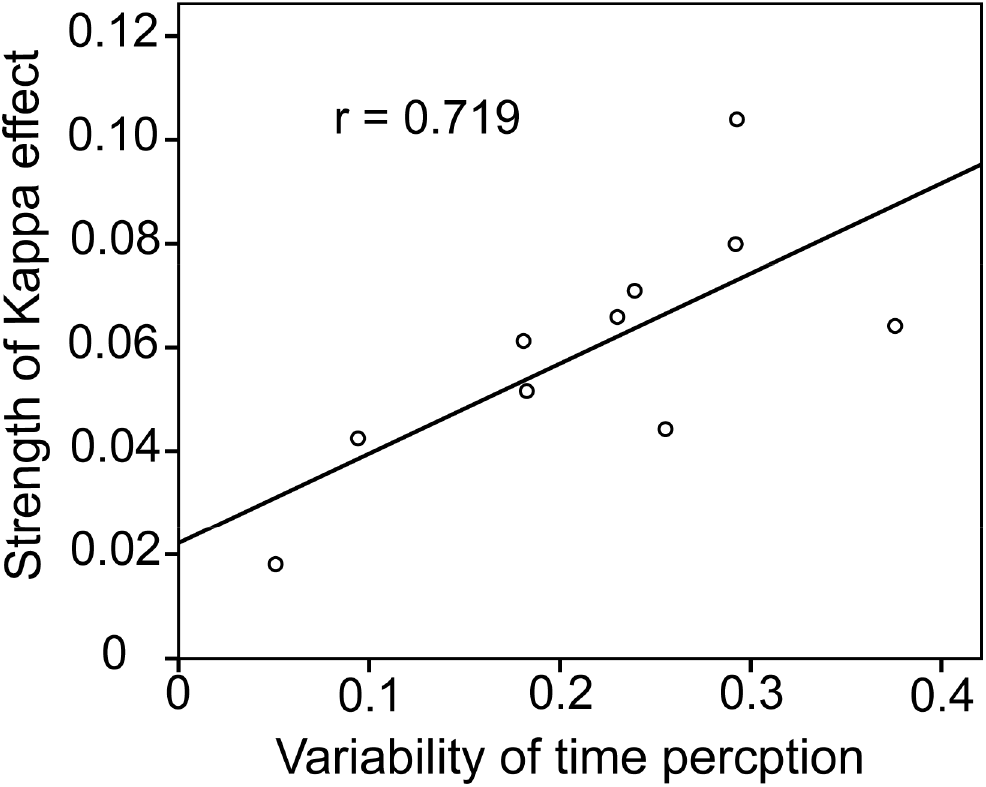
The correlation between variability of time perception (σsm) and the strength of the Kappa effect (κ = 1 − ωs).

We evaluated the goodness of model fitting by AIC index. A larger AIC means that the fitted model is less plausible. AIC Δ was obtained by subtracting AIC of the LCM from that of the classical model for each participant. AIC Δ ranged from 14 to 50. According to the levels of empirical support for model (0 < Δ < 2, substantial; 4 < Δ < 7, considerably less; Δ > 10, essentially none) [54], the LCM was superior to the classical model in the model fitting for all participants.

## Discussion

Here we used a time discrimination task to replicate the Kappa effect that the point of subjective equality (PSE) increases with the distance between two circles. We manipulated the vertical distance between circles and central fixation, and provided evidence that the tendency of deceleration of the Kappa effect is not driven by the variance of circle locations in Experiment 1 (Fig. 3). Then we manipulated distances between sample circles from 1° to 32°, and compared the fits of the classical model and the LCM for the tendency of deceleration in Experiment 2 (Fig. 4). AIC index provided quantitative evidence that the LCM fits data better than the classical model.

We found that the main effect of vertical distance was not significant, which indicates that the spatial variance of the circle locations does not influence the visual Kappa effect. Our result is not consistent with the prediction of the slowness model. The slowness model predicts that a given distance with a larger spatial variance of stimuli locations will be perceived as shorter[27], and that a shorter distance leads to a shorter duration perception[20,25]. This contradiction may be because of different characteristics of spatial processing in the visual and tactile modalities. Our ability to localize touch on body parts is often inaccurate. Take the cutaneous rabbit effect as an example, subjects can perceive a series of taps on the skin that does not exist[55,56]. However, our visual localizing is very precise. People can hit rapidly moving balls with amazing precision[57]. As predicted by the Bayesian observer model, tactical spatial processing is more inaccurate, thus the tactile processing is easier to be interfered by expectation[27]. Therefore, the slowness model is not suitable for explaining the visual Kappa effect.

Both the classical model and the LCM predict that the estimated time increases as an distance between two circles increases. However, the LCM is superior to the classical model in data fitting. The classical model predicts a linear relation between estimated time and distance (Equation 1 and 6). According to the LCM, the expected time is a ratio of a given distance and the constant speed, that is logarithmically transformed into internal expected time. The LCM predicts a logarithmic relation between the internal estimated time and the given distance (Equation 12) and a power relation between the PSE and the distance between two sample circles (Equation 18). The LCM replicated the tendency of deceleration of the Kappa effect (Fig. 4). The results suggest that the tendency of deceleration is driven by the Weber-Fechner law.

The key hypothesis of the LCM is a logarithmic internal representation of time. The logarithmic internal time hypothesis was deduced from the scalar variability of time perception, in which the standard deviation of timing increases linearly as the time interval increases. The scalar variability is consistent with the Weber-Fechner law which determines a logarithmic internal representation of time[34,35,44,45]. However, several studies reported that time estimation follows Stevens’ power law[58–60]. Previous studies revealed that the Bayesian model provides a direct link between Weber-Fechner and Stevens’ power law[31,37]. Our Bayesian model shows that the internal time is a logarithmic function of physical time and physical distance (Equation 7, 8, 12), which follows the Web-Fechner law. A motor response *t_e_*, that is an estimated time in the time reproduction task or verbal estimation task, is a power function of physical time and physical distance (Equation 14), and the PSE is a power function of physical distance (Equation 18), which follows the Stevens’ power law. Therefore, our model provides an integrated theoretical framework to consider the logarithmic time representation of power time estimation.

The speed prior is the basic hypothesis of the three Bayesian models. The slowness model assumes a low speed prior that objects tend to move at a low speed with a mean of zero[22–24]. The classical model and the LCM assume a constant speed prior that objects tend to move at a constant speed[19]. We found that the estimated constant speed of the classical model (0.25 °/s) is close to the absolute threshold of speed for older people with a mean age of 62 (0.12 °/s), and the estimated constant speed of the LCM (0.09 °/s) is equal to the absolute threshold of speed for younger people with a mean age of 23[61]. Thus our results confirmed a previous finding that the constant speed is slow[25]. These results provide conclusive evidence that the Kappa effect is driven by slow speeds.

We defined *κ* as an index of the strength of the Kappa effect, and found *κ* correlated with variability of time perception positively. To the best of our knowledge, we firstly reported the individual difference of the Kappa effect, in which individual with a more precise time perception would have a weaker Kappa effect. This result is consistent with a widely accepted notion of Bayesian theory that humans increasingly rely on prior knowledge as the uncertainty of a task increases[23,62–64]. Spatiotemporal interferences were especially correlated to depend on the variability of perceived dimensions, which is consistent with the prediction of Bayesian theory[65,66]. The above studies only provide qualitative evidence (interference vs. no interference), however, our study not only provides a theoretical function between the strength of the Kappa effect and variability of time perception 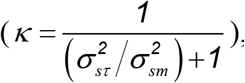 but also provides quantitative empirical evidence (Fig. 5).

The Bayesian model provides a new perspective to consider the cross-dimensional interferences. Our Bayesian model is useful in examining the Kappa effect with the constant speed (or slow speed) hypothesis. This Bayesian framework may be applied to explain other types of cross-dimensional interferences with appropriate assumptions. Taking interferences between time and the length of a line as an example, it is reasonable to consider that people have a prior that “a longer length takes a longer time to travel”[2,66]. When participants are required to estimate the presentation time of a line, we can assume an implicit motion along the line. The expected time is the ratio of the line length and a constant speed (*E*(*t*)=*l*/*v_0_*). This idea is consistent with a hypothesis that magnitudes co-vary across dimensions[66]. The other details of the model are identical with the present study. We assume that the implicit time estimation of expected motion is driven by the slow speed prior, and predict that the estimated constant speed is close to the absolute threshold of speed.

## Conclusion

We performed two experiments to find out why spatiotemporal interferences are accompanied by a tendency of deceleration. In Experiment 1, we found the variance of stimuli locations did not effect the estimated time, which suggests that the slowness model, originating from tactile neural activities, is not suitable for explaining the visual Kappa effect. In Experiment 2, we found that the Bayesian model on logarithmic scales made better behavioral predictions than the linear model, and provided a theoretical framework to integrate the logarithmic time representation with power time estimation. The estimated constant speed was close to the absolute threshold of speed, which confirms a previous finding that the Kappa effect is driven by slow speeds. Based on the logarithmic Bayesian model, we defined κ as an index of the strength of the Kappa effect, and that κ correlated with variability of time perception positively. Our findings suggest that the tendency of deceleration in the spatiotemporal interferences is driven by the Weber-Fechner law. This Bayesian framework aids in explaining the Kappa effect, and may be applied in the field of time perception and other types of cross-dimensional interferences with appropriate assumptions in future work.

## Acknowledgments

This study was supported by Humanities and Social Science Youth Foundation of Ministry of Education of China (Grant No. 19YJC190002) and Fundamental Research Funds for the Central Universities (Grant No. SWU1909444).

